# DNAHX: a novel, non-motile dynein heavy chain subfamily, identified by cryo-EM endogenously

**DOI:** 10.1101/2025.01.18.633724

**Authors:** Pengxin Chai, Diego Suchenski Loustaunau, Wan Zheng, Jun Yang, Kai Zhang

## Abstract

Ciliogenesis and cilia motility rely on the coordinated actions of diverse dyneins, yet the complexity of these motor proteins in cilia has posed challenges for understanding their specific roles. Traditional evolutionary analyses often overlook key family members due to technical limitations. Here, we present a cryo-EM-based, bottom-up approach for large-scale, *de novo* protein identification and functional prediction of endogenous axonemal dynein complexes. This approach led to the identification of a novel dynein heavy chain subfamily (XP_041462850), designated as DNAHX, from sea urchin sperm. Phylogenetic analysis indicates that DNAHX branches from the outer-arm dynein alpha chain during evolution and is found in specific animal lineages with external fertilization. DNAHX contains multiple insertions throughout the protein, locking DNAHX permanently in a pre-powerstroke state. The AAA1 site exhibits poor conservation of essential ATPase motifs, consistent with DNAHX’s non-motile nature. DNAHX also forms a heterodimeric dynein complex, which we named dynein-X, with another dynein heavy chain and accessory chains. Furthermore, a subset of dynein-X displays an autoinhibited phi particle conformation, potentially facilitating the intraflagellar transport of axonemal dyneins. Our discovery of the novel, non-motile dynein heavy chain and the dynein-X complex provides valuable insights into the evolution of dyneins and potentially their diverse cellular functions.

## Introduction

Dyneins are evolutionarily conserved cytoskeletal motor proteins that move toward the minus-end of microtubules (MTs) [1]. Based on their cellular roles and locations, dyneins are classified into cytoplasmic and axonemal groups [2]. Cytoplasmic dyneins enable retrograde transport of various cellular cargoes along MTs and are subdivided into dynein-1, for intracellular transport in cytosol [3], and dynein-2, for intraflagellar transport within cilia [4]. Axonemal dyneins, consisting of outer-arm dynein (OAD) and inner-arm dynein (IAD), are anchored on axonemal doublet MTs (DMTs) [5, 6] and facilitate the coordinated beating of cilia [7, 8, 9, 10]. Given their essential functions, dynein defects are linked to neurodegenerative diseases [11] and primary ciliary dyskinesia [12].

All dynein complexes share a core architecture with one or more dynein heavy chains (DHC) and several accessory proteins [13, 14]. Each DHC contains an N-terminal tail domain, responsible for interacting with accessory proteins and cofactors, and a C-terminal motor domain, from the AAA+ protein family (ATPases Associated with Diverse Cellular Activities), which drives MT-based motility [15, 16, 17]. The extensive sequence information from DHC (∼4,500 amino acids, ∼500 kDa) supports phylogenetic analysis of DHC genes, revealing distinct dynein subfamilies correlating to the specific dynein types [18, 19, 1, 20, 21]. For example, dynein-1 and dynein-2 DHC subfamilies are closely related, consistent with their shared cargo transport roles. And the OAD-*α, β* and *γ* HC subfamilies together constitute OAD complexes.

In addition to these canonical DHC genes, phylogenetic analyses have revealed numerous non-canonical or “orphan” DHC genes across species [22]. The most well-known is DNHD1, the only “orphan” DHC encoded in human genome [2]. Previously thought to be a non-functional relic from early DHC gene duplication [23], recent evidence indicates that DNHD1 localizes within axonemes, where it plays a critical role during ciliogenesis in nodal cilia [24] and sperm flagella [25, 26]. This suggests that DNHD1 and other orphan DHC genes may have evolved specific roles related to ciliary and flagellar assembly and regulation. However, dynein is typically a complex containing both heavy chains and accessory chains, and little is known about the structures and mechanism of the whole complex that incorporate these orphan DHC genes.

Unlike canonical DHC genes with well-defined cellular roles and locations, functional interpretation of orphan DHC genes remains challenging. Firstly, traditional phylogenetic analyses provide only sequence information and cannot associate these orphan DHCs with structural information of whole dynein complex. Secondly, because orphan DHCs likely participate in ciliary assembly and regulation, the complexity of dynein types involved in these processes obscures their precise roles [4]. To overcome the challenges and expand our understanding of orphan DHCs and dynein complexes within cilia, we developed a bottom-up, cryo-EM-based approach for large-scale, *de novo* protein identification of native axonemal dyneins, by expanding our recent approach to analyzing dynamic structures of the full-length human dynein-1 actively engaged in its mechanochemical cycle [27]. Applying this technique to multiple dyneins from sea urchin sperm, co-exising in the same sample in the presence of ATP, we resolved both compositional and conformation heterogeneity of axonemal dyneins simultaneously, leading to the *de novo* identification of a new DHC gene, XP_041462850, which we named DNAHX. Phylogenetic analysis indicates that DNAHX diverged from the OAD-*α* HC. Structural analysis revealed that DNAHX contains multiple insertions and lacks conservation of key ATPase motifs at its active site, suggesting a loss of microtubule-based motility. Intriguingly, DNAHX assembles into a heterodimeric dynein complex, termed dynein-X, alongside another DHC, which is not DNAHX itself, and accessory chains. A subset of dynein-X complexes adopts an autoinhibited phi particle conformation, suggesting a potential role in axonemal dynein transport and regulation. These findings uncover previously uncharacterized structural and functional diversity in the dynein family, providing a framework to further explore the evolution and specialization of many dynein-mediated cellular processes.

## Results

### Workflow of Protein Identification of Native Dynein Complexes using Cryo-EM

We have recently demonstrated that cryo-EM is capable of resolving fine details of conformational changes while dynein is actively engaged in its mechanochemical cycle [27]. To further test whether this dynamic structural analysis approach can be extended to study more complex systems with multiple dynein types with various compositions and conformations, we selected the green sea urchin (*Lytechinus variegatus*) as our source to isolate two-headed axonemal dyneins (**Fig. 1a**). Negative-staining microscopy followed by 2D class classification analysis confirmed that the final purified sample predominantly consisted of two-headed dynein complexes (**Fig. 1b**). Given that OAD comprises different isoforms in higher eukaryotes—such as OAD-*α* heavy chains (HC: DNAH5 and DNAH8) and OAD-*β* HC (DNAH9, DNAH11, and DNAH17) [28]—we employed mass spectrometry (MS) to analyze the composition of the isolated dynein complexes and quantify the relative abundance of each dynein heavy chain (**Fig. 1c**).

**Figure 1:**
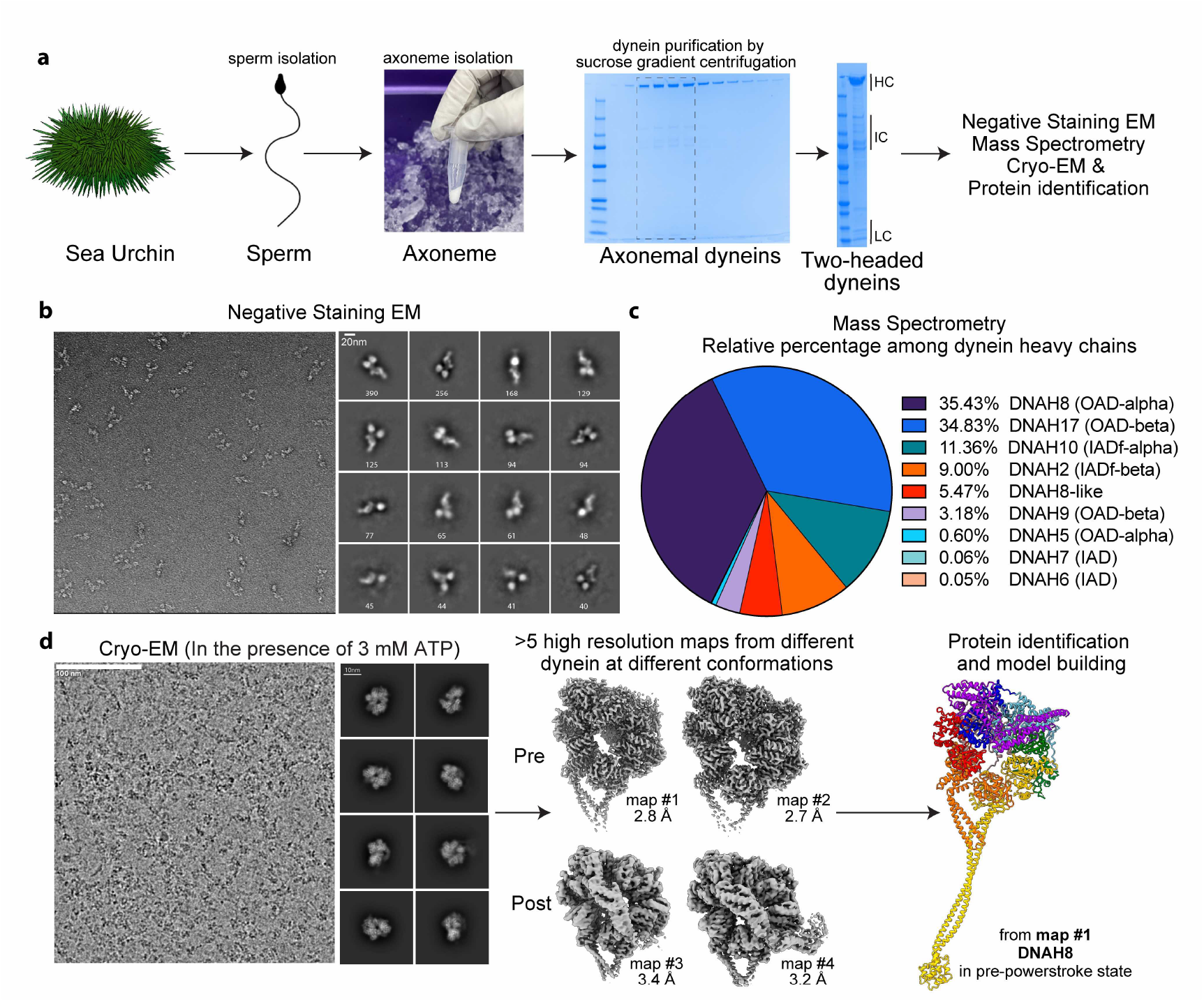
Workflow for protein identification of native dynein complexes using cryo-EM. **(a)** Flowchart outlining the isolation of sea urchin sperm, preparation of axonemes, and purification of two-headed dynein complexes. **(b)** Representative negative staining electron micrograph of the purified dynein complex. The 2D class averages, predominantly showing two-headed dynein, are displayed on the right. **(c)** Mass spectrometry-based quantification of dynein heavy chains from the purified sample. **(d)** Cryo-EM analysis and protein identification of different dynein heavy chains at various stages of the powerstroke cycle.

Our MS results reveal that isolated dynein sample mainly consists of two OADs, DNAH8 (35.43%) and DNAH17 (34.83%), as well as two minor OAD HCs, DNAH5 (0.6%) and DNAH9 (3.18%), aligning with prior findings that DNAH8 and DNAH17 are preferentially expressed in sperm cells [28]. Additionally, inner-arm dynein-f, comprising DNAH10 (11.36%) and DNAH2 (9.00%), was co-isolated, consistent with known cryo-ET and cryo-EM structure of inner-arm dynein-f as a two-headed dynein [29, 9]. Alongside these dynein heavy chains, MS analysis also detected other DHCs, including a DNAH8-like protein and two single-headed inner-arm dyneins (IAD) identified as DNAH7 and DNAH6.

To gain high-resolution structural insights into the dynein complexes in reaction, we prepared cryo-EM samples in the presence of 3 mM ATP (**Fig. 1d**), similar to the approach we previously described [27]. The 2D class averages revealed key motor domain features, such as AAA+ domains and the linker region. We then performed *in silico* 3D classification analysis followed by focused refinement to obtain high-resolution cryo-EM reconstructions (**Supplementary Fig. 1a, b**). Building upon our recent study of cytoplasmic dynein-1 alone in the presence of ATP [27], the current work extends beyond that simpler system to encompass a broader array of axonemal dynein heavy chains, resolving their structures at multiple states (both pre- and post-powerstroke) all together.

To identify the corresponding DHC genes, we employed AI-based automatic model-building tools [30, 31], which typically generate models with several polypeptide chains for each map. The sequence of the longest polypeptide chain (∼ 1000 amino acids) was then used for BLASTP searches against the NCBI *Lytechinus variegatus* protein database. Owing to the high resolution of the cryo-EM maps (better than 3.5 Å) and the large molecular weight of the dynein motor domain (>350 kDa), this approach was sensitive enough to distinguish between different HC genes, including HC isoforms. For instance, for map #1 with a resolution of 2.7 Å, the top BLASTP hit was DNAH8 with 77.05% identity, while the second hit was DNAH5 with 60.99% identity (**Supplementary Fig 2a, b**). Following protein identification, atomic models were constructed for each map.

### Identification and Phylogenetic Analysis of a Novel Dynein Heavy Chain: DNAHX

From our cryo-EM maps and atomic models, we noticed a DHC gene (NCBI Reference Sequence: XP_041462850), annotated as a DNAH8-like protein, with an unusually long sequence of 5,272 amino acids (**Fig. 2a**). Given that typical dynein heavy chains are approximately 4,500 amino acids long, we speculated that this could represent a novel type of DHC. To investigate this possibility, we constructed a phylogenetic tree using this gene alongside known DHC genes from several species previously utilized for DHC classification [32] (e.g., *Homo sapiens, Lytechinus variegatus, Ashbya gossypii, Caenorhabditis elegans, Chlamydomonas reinhardtii, Danio rerio, Drosophila melanogaster, Kluyveromyces lactis*, and *Tetrahymena thermophila*). Our analysis revealed that XP_041462850 formed a distinct branch, with the closest related gene being the OAD-*α* HC (Fig. 2b). Based on this observation, we designated the gene “DNAHX,” where “X” symbolizes the unknown nature of this novel DHC.

**Figure 2:**
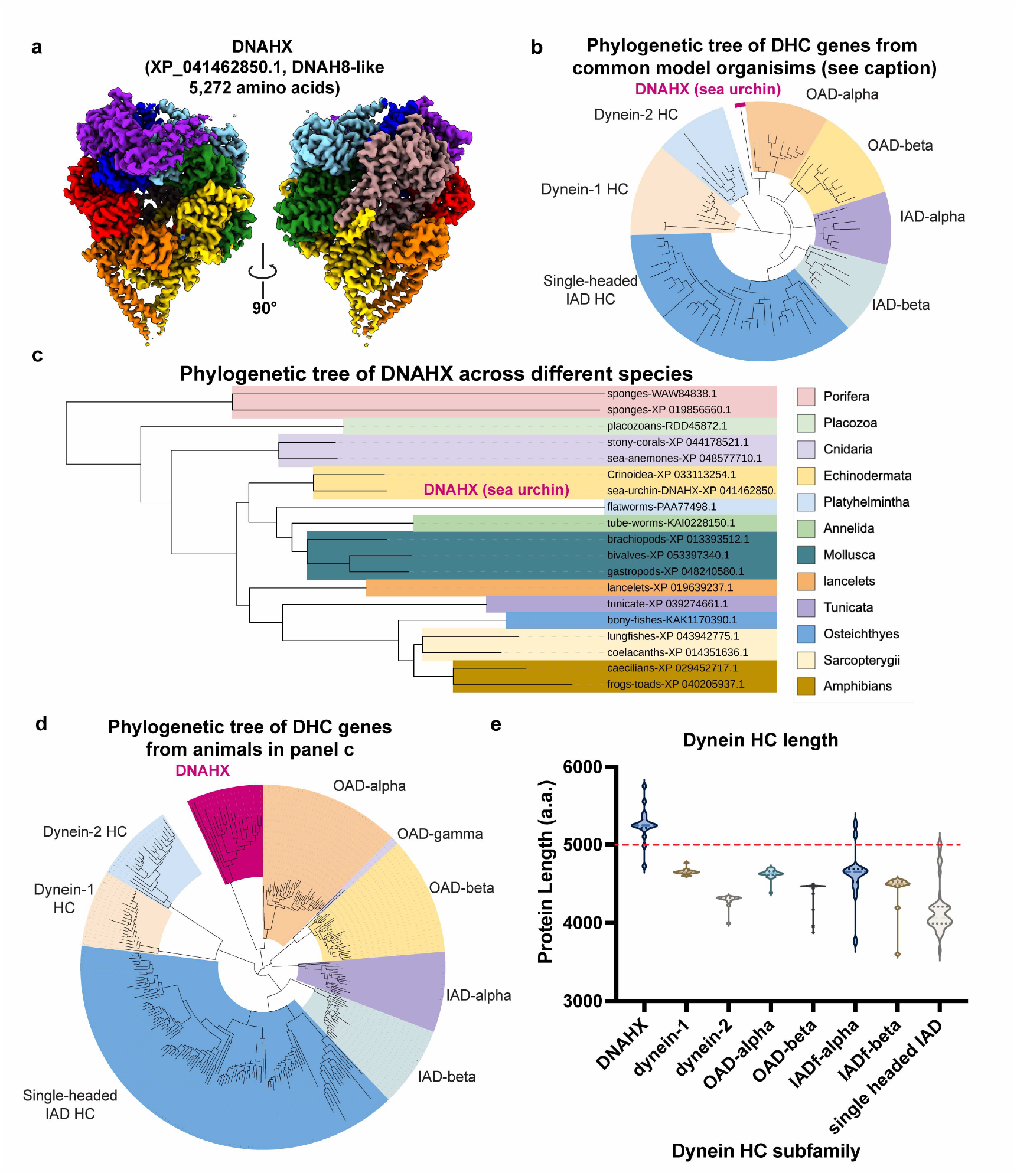
DNAHX is a novel dynein heavy chain subfamily characterized by its extended protein length. **(a)** Cryo-EM density map of the DNAHX protein (XP_041462850) with unusually long sequence with 5,272 amino acids. **(b)** Phylogenetic classification of DNAHX with known dynein heavy chains from various species, including *H. sapiens, L. variegatus, A. gossypii, C. elegans, C. reinhardtii, D. rerio, D. melanogaster, K. lactis, and T. thermophila*. **(c)** Phylogenetic tree depicting the evolutionary relationships of DNAHX across different species. **(d)** Detailed phylogenetic classification of DNAHX alongside known dynein heavy chains from species that possess DNAHX. **(e)** Comparative analysis of protein length across different dynein heavy chain families, emphasizing the extended length of DNAHX relative to other dyneins.

Interestingly, DNAHX was not found in the aforementioned species, with the exception of sea urchin, suggesting that DNAHX may be a species-specific DHC gene. To further investigate the homologs of DNAHX, we conducted a BLASTP search against the entire NCBI database. To confirm the top hit as a true homolog of DNAHX and not another axonemal DHC, we performed a reverse BLASTP search of the hit against the *Lytechinus variegatus* database. This approach enabled us to construct a phylogenetic tree of DNAHX across various species (**Fig. 2c**). The analysis revealed that DNAHX was absent in protists, such as *Tetrahymena* and *Chlamydomonas*, which are model organisms for motile cilia research. Additionally, DNAHX was found only in a subset of animals, ranging from primitive multicellular organisms like sponges to vertebrate amphibians like frogs. Notably, DNAHX was absent in reptiles and mammals. Since DNAHX and its homologs are mainly from the organisms living in water, their restricted presence might be related to the mode of fertilization, as those species with DNAHX typically release sperm into water for external fertilization.

To further validate the DNAHX being a novel DHC subfamily, we constructed another phylogenetic tree using DNAHX and known DHC genes from animals that have DNAHX. As anticipated, DNAHX emerged as a distinct branch, likely originating as a duplication of the OAD-*α* HC during evolution (**Fig. 2d**). We also analyzed the protein length across various DHC genes and confirmed that the unusually long protein length is a distinctive feature of DNAHX (**Fig. 2e**).

### DNAHX Shows Extra Insertions at Multiple Sites

To investigate the reasons behind the longer protein length of DNAHX compared to other DHCs, we performed sequence and structural comparisons between sea urchin DNAHX and its closest relative, DNAH8. Using the same dataset, we resolved the motor domains of both DHCs at high resolution (**Supplementary Fig. 1**,**2**). To construct full-length models of these DHCs, we used AlphaFold2 [33, 34] for structural predictions of the flexible regions not resolved in the cryo-EM map, such as the tail domain and stalk-MTBD region (**Supplementary Fig. 3a, b**). This analysis revealed that DNAHX contains multiple insertions distributed across the entire heavy chain. Many of these insertions are disordered and flexible, including a prominent insertion in the tail region (**Supplementary Fig. 3b)**, explaining the unusually long protein length of DNAHX.

We then focused on the motor domain (**Fig. 3a, b**), a critical region for dynein function and ATPase activity [35, 36]. Previous studies have shown that dynein motor domains often carry insertions, such as those located between AAA large/small (L/S) subunits and within AAA domains. For instance, the H2 insert at the AAA2L domain in dynein has been implicated in regulatory functions [37]. In DNAHX’s motor domain, we identified several additional insertions compared to DNAH8, particularly in the stalk and MTBD regions and within AAA4S. We hypothesize that these insertions may influence dynein function. To explore this further, we conducted a detailed analysis of each insertion, which is described below.

**Figure 3:**
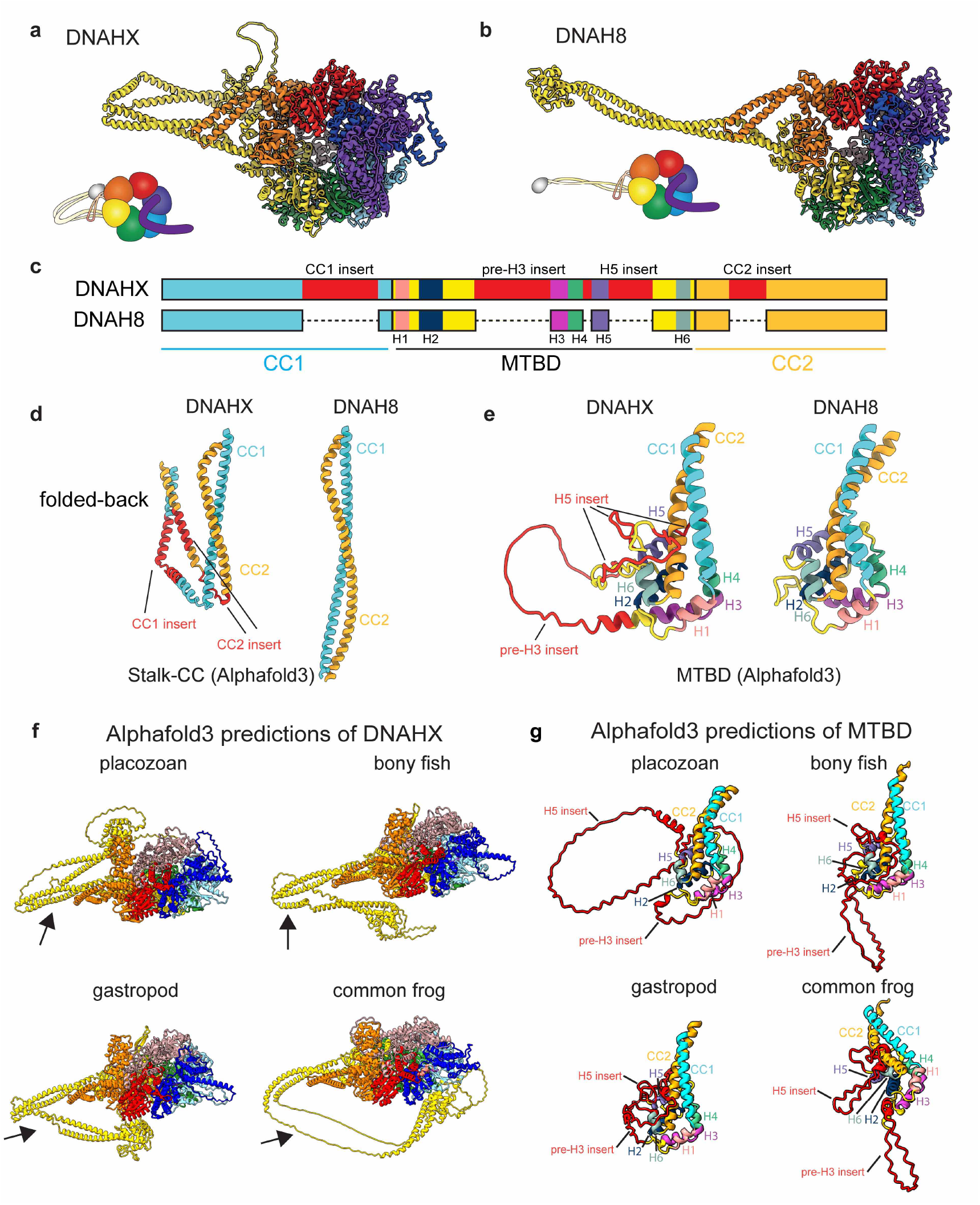
Insertions in the stalk-MTBD region disrupt microtubule-binding capacity. **(a, b)** Cartoon models of DNAHX and its closest relative, DNAH8, with individual domains colored. Models were constructed using a combination of cryo-EM density maps and Alphafold2 predictions. **(c)** Domain architecture of the stalk-MTBD region in DNAHX and DNAH8. Insertions in DNAHX are highlighted in red. **(d)** Structural comparison of the coiled-coil stalk between DNAHX and DNAH8. Insertions in DNAHX cause a “folded-back” conformation of the stalk. **(e)** Structural comparison of the microtubule-binding domain (MTBD) between DNAHX and DNAH8. **(f)** Alphafold3 predictions of the DNAHX motor domain from four additional species, showing similar structural features. **(g)** Detailed view of the MTBD in DNAHX from different species, highlighting the pre-H3 and H5 insertions of varying lengths.

### Insertions at Stalk-MTBD Disrupts the Microtubule-Binding Capacity

We find that the stalk-MTBD region of DNAHX contains several insertions (**Fig. 3c**). While the stalk of DNAH8 exhibits a canonical coiled-coil conformation, as predicted by AlphaFold2, the stalk of DNAHX displayed a novel conformation with the MTBD folded back toward the AAA ring. This altered conformation is due to insertions in the coiled-coil-1 (CC1) and CC2 regions, which disrupt the hydrophobic patches of the coiled-coil, causing additional kinks in the helices (**Fig. 3d**). Additionally, the MTBD of DNAHX contains disordered insertions, specifically before helix-3 (pre-H3) and within the helix-5 (H5) region (**Fig. 3e**), which may affect the MT-binding ability.

To determine whether these insertions in the stalk-MTBD region are a common feature of the DNAHX subfamily, we performed AlphaFold3 [33] predictions of motor domains from DNAHX orthologs in four diverse species: placozoan, bony fish, gastropods, and the common frog. We found that the stalk regions in all four structures contain insertions of varying lengths, which modify the folding of the coiled-coil (**Fig. 3f**). Similarly, the MTBDs from these species also exhibit disordered pre-H3 and H5 insertions of different lengths (**Fig. 3g**), suggesting that these insertions do not serve a conserved functional role. This contrasts with a previously studied “flap” insertion at MTBD that binds microtubules with a conserved length and folding [38, 39].

In conclusion, the insertions in the stalk-MTBD region likely disrupt the microtubule-binding ability of DNAHX.

### Insertions at AAA4 Small Domain Locks DNAHX in Pre-Powerstroke State

In addition to the disordered or flexible insertions found in DNAHX, we identified a well-folded tertiary structure insertion in the AAA4 small domain (AAA4S), which features a *β*-hairpin motif and an *α*-helix (**Fig. 4a**). This insertion occurs between helices H6 and H7 in AAA4S and is classified as a presensor II insert (PS2i). While PS2i has been observed in other AAA+ proteins such as MCM and RavA [40], it has not been previously found in dynein AAA+ domains. We noted several differences between the PS2i in DNAHX and those previously observed. First, canonical PS2i forms a long *α*-helix, whereas PS2i in DNAHX includes an additional *β*-hairpin motif, typically found in pre-sensor 1 and helix-2 inserts. Additionally, while PS2i generally functions to reposition the small subdomain, no significant rearrangement of AAA4S was observed in DNAHX, suggesting that this PS2i may have a different functional role.

**Figure 4:**
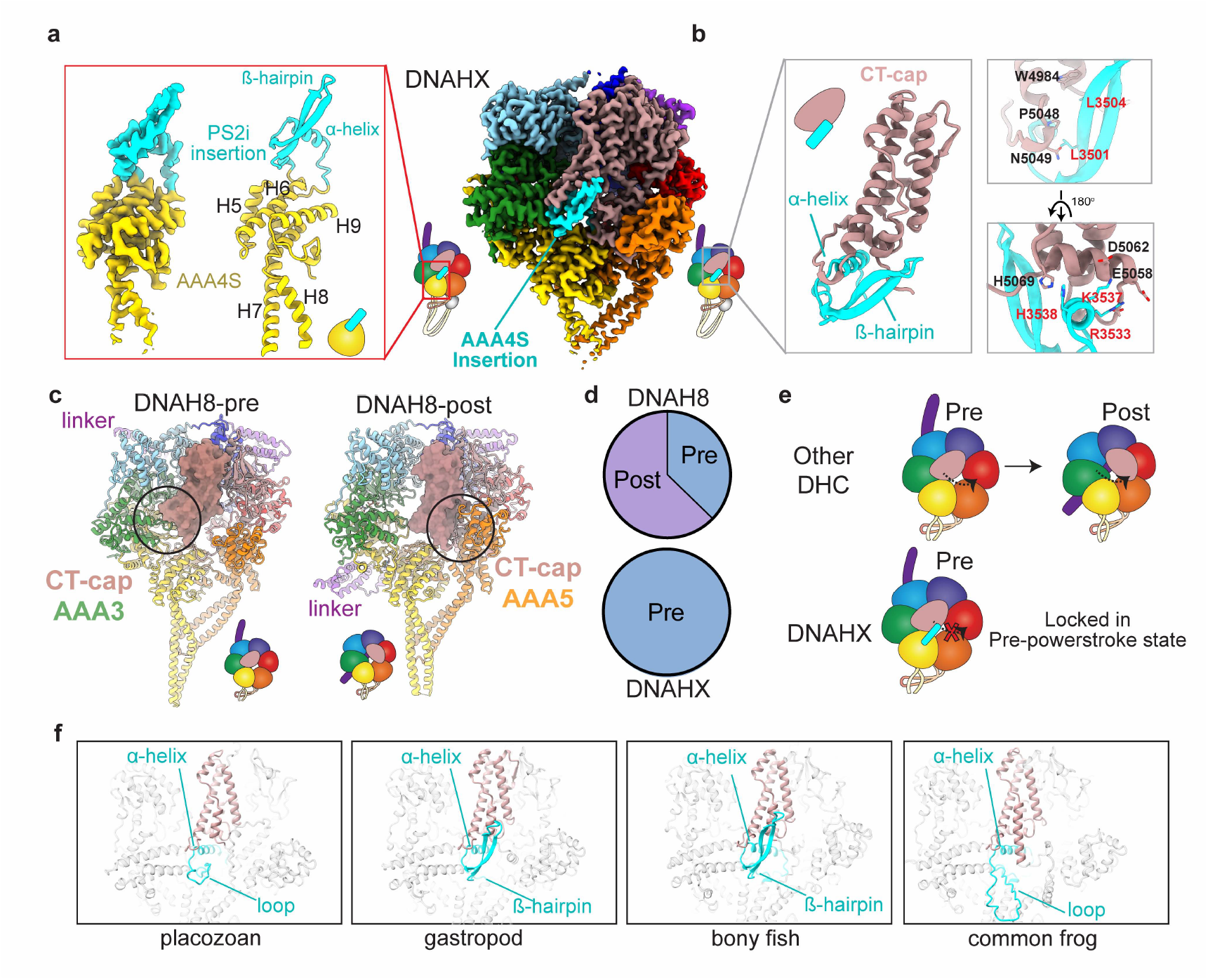
Insertions in the AAA4 small domain lock DNAHX in a pre-powerstroke state. **(a)** Cryo-EM density map and cartoon model of the AAA4S domain with the PS2i insertion. The right panel shows the cryo-EM density map of the DNAHX motor domain. **(b)** Cartoon representation of the molecular interactions between the PS2i and the CT-cap. The residues involved in the interaction are highlighted as sticks. **(c)** Structural transition of the CT-cap between pre-powerstroke and post-powerstroke states. **(d)** Pie chart depicting the distribution of pre-powerstroke and post-powerstroke states in DNAH8 and DNAHX, derived from the same cryo-EM dataset. **(e)** Model illustrating how the PS2i insertion restricts the CT-cap swing, locking DNAHX in the pre-powerstroke state. **(f)** Conserved PS2i found in other members of DNAHX. Note that the *β*-hairpin motif in some DNHAX is not stale, but the *α*-helix is all conserved and is predicted to interact with CT-cap.

Notably, PS2i specifically interacts with the C-terminal cap (CT-cap) via the *β*-hairpin motif and *α*-helix, forming a series of hydrophobic and salt-bridge interactions (**Fig. 4b**). The CT-cap is known to modulate the mechanochemical cycle of dynein and has been shown to swing from AAA3 to AAA5 as dynein transitions between pre-powerstroke and post-powerstroke states [27, 41, 42]. In agreement with this, we observed that the CT-cap in DNAH8 could shift its docking sites between AAA3 (pre-powerstroke) and AAA5 (post-powerstroke) in our cryo-EM analysis (**Fig. 4c, d**). By contrast, no post-powerstroke state of DNAHX was detected in the same dataset (**Fig. 4d**), suggesting that the PS2i at AAA4S may prevent the CT-cap from swinging and regulate DNAHX’s mechanochemical cycle (**Fig. 4e**). Furthermore, we identified PS2i in other members of the DNAHX family, and structural predictions indicate similar interactions with the CT-cap, potentially implying a conserved regulatory mechanism (**Fig. 4f**).

### Non-conserved AAA1 Site of DNAHX is Locked in ATP-binding State

Our cryo-EM results reveal that DNAHX is likely locked in a pre-powerstroke state while other axonemal dynein complexes show both pre- and post-powerstroke in the presence of ATP, suggesting that dynein-X likely functions as a non-motile motor. This drove us to further investigate the structural details of its AAA1 pocket, the primary ATP hydrolysis site [35]. Sequence alignment among DHCs from sea urchins reveals that the AAA1 site in DNAHX is distinct from other DHCs, particularly in several key motifs including Walker-A, Sensor-I, and Sensor-II loops (**Fig. 5a**).

**Figure 5:**
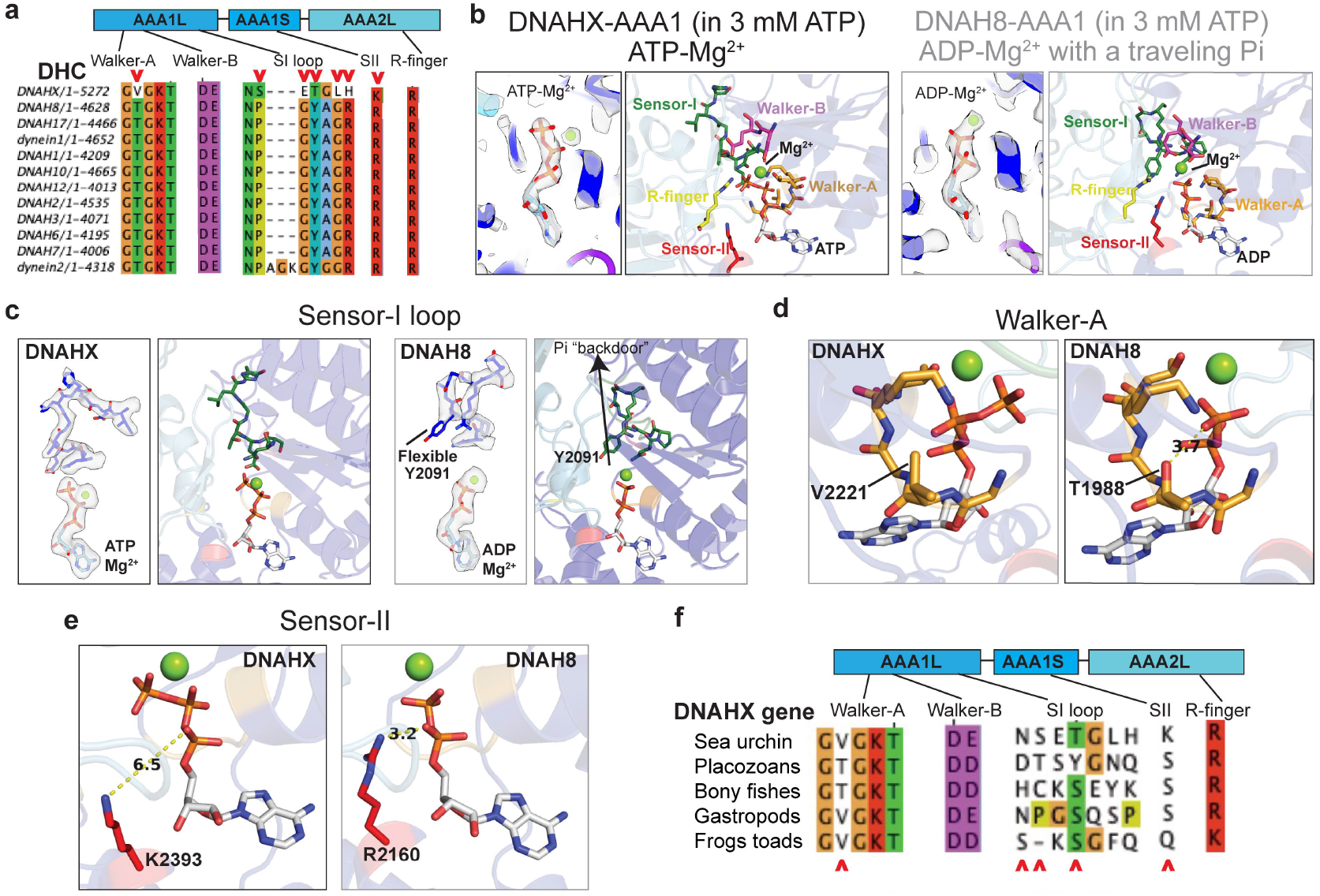
Variations in key motifs lock DNAHX-AAA1 in an ATP-bound, pre-powerstroke state. **(a)** Sequence alignment of the AAA1 site in DNAHX compared to other dynein heavy chain genes in sea urchin, highlighting variations in critical motifs such as Walker-A, Sensor-I, and Sensor-II loops. **(b)** Comparison of nucleotide states at the AAA1 site between DNAHX and DNAH8. DNAHX shows ATP-binding at the AAA1 site, while DNAH8 exhibits ADP-Mg^2+^ density in cryo-EM maps. The cartoon models of active site are shown. **(c)** Structural details of the Sensor-I loop in DNAH8 and DNAHX. DNAH8 retains a flexible tyrosine (Y2091) essential for the “molecular backdoor” function, while DNAHX has mutations that stabilize the loop, indicating the loss of this function. **(d)** Structural comparison of the Walker-A motif in DNAH8 and DNAHX. The threonine in DNAH8 is replaced by valine in DNAHX, disrupting hydrogen bonding with phosphate groups. **(e)** The arginine residue in the Sensor-II motif of DNAH8 is replaced by lysine in DNAHX, weakening interactions with phosphate groups. **(f)** Comparative analysis of AAA1 motifs across the DNAHX subfamily reveals that these motifs are not conserved.

We then compared the nucleotide states of AAA1 in DNAHX and DNAH8 in the pre-powerstroke conformation (**Fig. 5b**). Surprisingly, we find that DNAHX retains an ATP molecule at the AAA1 site, suggesting that it is permanently trapped in a pre-hydrolysis state, despite the presence of a Walker-B motif. By contrast, DNAH8 shows clear density for ADP-Mg^2+^ and a flexible Sensor-I loop in the presence of ATP, indicating its ATP hydrolytic activity (**Fig. 5c**). We previously have demonstrated that the Sensor-I loop in dynein-1 acts as a molecular backdoor”, mediating the phosphate release [27]. Specifically, the tyrosine residue (Y2022 in human dynein-1) within the Sensor-I loop undergoes a significant conformational change to gate the opening or closure of the backdoor. The observation of a similarly flexible Y2091 in DNAH8 supports this model. However, the Sensor-I loop in DNAHX largely differ other dyneins. Notably, the residues proline and tyrosine (P2089, Y2091 in DHAH8) that are conserved among other dyneins are replaced with serine and threonine (S2322, Y2324 in DHAHX), respectively. The Sensor-I loop in DNAHX is also stabilized as clearly visualized in the cryo-EM density, implying that its canonical function as a molecular backdoor for phosphate release has been lost during evolution.

In addition, we observed that the conserved threonine residue (T1988) in the Walker-A motif is replaced with valine (V2221) in DNAHX (**Fig. 5d**). This substitution disrupts the hydrogen bond between the hydroxyl group of threonine and phosphate groups, potentially hindering its ATP hydrolysis. Furthermore, the conserved arginine residue (R2160) in the Sensor-II motif is replaced with lysine (K2393) in DNAHX (**Fig. 5e**). While lysine maintains a positive charge, it does not form hydrogen bonds or electrostatic interactions with phosphate groups. In contrast, the arginine residue in DNAH8, with its more complex guanidinium group, directly interacts with phosphate groups, which can be critical for ATP hydrolysis.

The differences in these key motifs likely contribute to the observed ATP-binding state of AAA1 in DNAHX, further supporting our observation that DNAHX is locked in a pre-powerstroke conformation and the hypothesis that AAA1 no longer plays a role in dynein mechanochemical cycle and microtubule-based motility. To further confirm our hypothesis, we compare the AAA1 motifs across various DNAHX genes. Our analysis indicates that the motifs within DNAHX subfamily vary substantially but they are all distinct from these classical dynein motors known to date, suggesting that the DNAHX subfamily is non-motile as a whole, owing to their low conservation at those critical sites essential for ATP hydrolysis (**Fig. 5f**).

### DNAHX Forms a Phi Particle Conformation with Another Dynein Heavy Chain

To further understand the potential function of this non-motile DNAHX, we re-analyzed our cryo-EM data to examine whether DNAHX forms a complete dynein complex, typically as a dimer as our isolation strategy preferentially enriches two-headed dyneins. We re-extract the DNAHX particles using a larger box size, as the original size only focuses one motor domain (**Fig. 6a**). Reference-free 2D classification analysis reveals that most DNAHX particles (60%) appear as monomeric dynein, while 27% form a two-headed dynein complex in an open conformation. Intriguingly, 13% of DNAHX particles appear in form of phi particle dynein (**Fig. 6b**). In addition to the additional DHC, the 2D class average also contains the densities for accessory chains.

**Figure 6:**
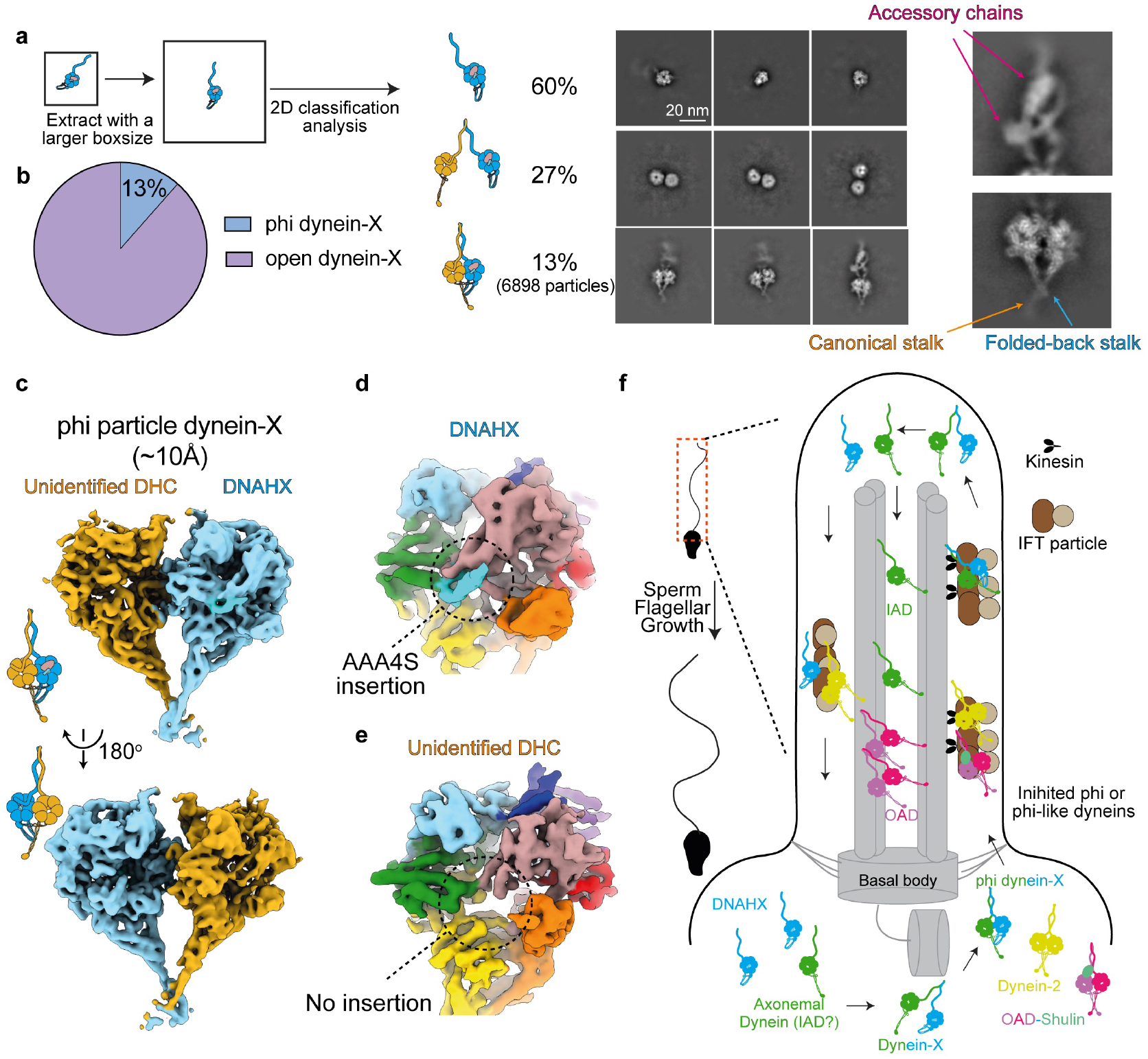
DNAHX forms a phi particle dynein complex with another DHC, named dynein-X. **(a)** Schematic of the workflow for re-extracting DNAHX particles using a larger box size to capture two-headed dynein species. The original box size only covered one motor domain. Reference-free 2D classification analysis of DNAHX particles. 60% of particles were monomeric dynein, 27% formed a two-headed dynein complex in an open conformation, and 13% formed a phi particle dynein. Arrows highlight two different stalk conformations. Representative 2D class averages are shown. **(b)** Pie chart showing the distribution of phi particle dynein-X and open dynein-X. **(c)** 3D reconstruction of phi particle dynein at 10Å resolution. **(d, e)** Close-up view of the two motor domains from phi particle dynein, showing that the second motor domain is not DNAHX, as it lacks the characteristic insertion at AAA4S. **(f)** Proposed model for DNAHX function in sperm cells. DNAHX, always locked in a pre-powerstroke state, serves as a template to form a phi particle with other axonemal dyneins, most likely single-headed IAD. This interaction inhibits IAD activity in the cytosol and facilitates its transport into growing cilia.

In the stalk-stalk dimerization region of the phi particle, one motor domain displayed a canonical extended stalk conformation, while the other motor domain lacks visible density beyond the stalk-stalk crossing region, indicating this phi dynein contains two different DHCs. The motor domain with the abnormal stalk conformation perfectly matches structural features of DNAHX, predicted to adopt a “folded-back” stalk conformation (as depicted in **Fig. 3**). To further verify this is DNAHX and reveal what the other DHC is, we further resolved a 3D reconstruction of this phi dynein at an overall resolution of 8Å (**Fig. 6c**). Our cryo-EM map clearly shows that the second motor domain is not DNAHX, as it lacks the characteristic insertion at AAA4S in DNAHX (**Fig. 6d, e**). We then named the dynein complex consisting of DHNAX, the other unidentified DHC and accessory chains as “dynein-X”.

The limited resolution of the cryo-EM map, due to the low number of particles, prevented definitive identification of the second DHC in the dynein-X complex. Our mass spectrometry data (**Fig. 1c**) suggest that the second motor domain may belong to one of the single-headed inner arm dyneins, such as DNAH6 or DNAH7. However, the possibility that DNAHX forms a dimer with two-headed OAD-HC or IADf-HC cannot be excluded.

To evaluate the candidates for the unknown DHC, we fitted AlphaFold-predicted motor domains in the pre-powerstroke state from various DHCs into the cryo-EM density map and calculated the correlation values (**Supplementary Fig. 4**). This analysis revealed that, apart from DNAHX, which exhibited a low correlation value of 0.76, all other fitted motor domains showed correlation values close to 0.80. These results indicate that multiple DHCs, including DNAH6, DNAH7, and others, could potentially form phi particle with DNAHX in the dynein-X complex.

The self-assembly of a heterodimeric dynein into a phi particle complex is a relatively rare phenomenon. Dynein-1 and dynein-2 heavy chains exclusively form homo-dimeric phi particles [43, 44]. Previously reported three-headed and two-headed OADs require additional cofactors, such as Shulin or DNAAF9, to adopt phi or phi like conformations [45, 46]. Only recently, the inner-arm dynein-f was also shown to adopt a phi conformation [47]. Our discovery of a phi particle dynein-X complex comprising DNAHX and another DHC strongly suggests that the phi particle conformation may represent a universal feature of two-headed dynein complexes, regardless of whether the dynein heavy chains are homodimeric or heterodimeric.

## Discussion

### A Novel Bottom-up Approach to Probe the Structure and Function of Axonemal Dyneins

In this study, we present a cryo-EM-based approach that directly probes the structures of a mixture of axonemal dyneins isolated from natural sources (**Fig. 1**). Using this method on sea urchin sperm, we are able to directly resolve high-resolution cryo-EM reconstructions of the dynein motor domain in multiple conformational states from different DHCs **(Fig. 1, 2**). Furthermore, this approach allows *de novo* identification of a new dynein family named DNAHX. Phylogenetic analysis and structural comparisons reveal this is a novel non-motile DHC, expanding our understanding of dynein diversity and function (**Fig. 2–6**). Our approach also holds the potential for studying various conformational states of dynein complexes and identifying novel forms, either as individual complexes, or together with novel cofactors, or by forming a complex like dynein-X, from many other possible sources such as mammalian tracheal cilia and sperm cells under various physiological, pathological and environmental conditions. With the fast-growing throughput of modern cryo-EM technology and high-performance computation, we foresee that similar approaches will substantially impact the dynein field and related research.

### DNAHX Sheds Light on Dynein and AAA+ Protein Evolution

DNAHX is identified as a non-motile DHC within the dynein family, offering valuable insights into the evolution of dyneins. Dynein-X containing DNAHX and another identified DHC is likely non-motile as well. Our phylogenetic analysis indicates that DNAHX is absent in protists, suggesting it arose after the emergence of multicellular animals. DNAHX likely originated from a duplication event of the outer-arm dynein *α*-HC in early animal evolution, as it is present in basal animals like sponges and placozoans (**Fig. 2**). Unique insertions and modifications at DNAHX’s AAA1 site suggest that this dynein evolved specifically to become non-motile.

Among these evolutionary changes, the mutation at the primary site of AAA1 may represent the initial event driving DNAHX to become a non-motile dynein, resulting in an abnormal mechanochemical cycle. Subsequently, the insertion in the stalk-MTBD likely impaired DNAHX’s microtubule-binding ability. Furthermore, an insertion at AAA4S may have permanently locked DNAHX in a pre-powerstroke state. Alternatively, it is also possible that these changes occurred concurrently, collectively contributing to the loss of microtubule-based motility.

A novel *β*-hairpin motif followed by an *α*-helix insertion within DNAHX’s AAA4 domain further expands the functional repertoire of AAA+ protein insertions [40]. Unlike typical AAA+ insertions, which are usually either a *β*-hairpin or an *α*-helix, DNAHX’s AAA4 contains both. Moreover, while the canonical PS2i insert is a single *α*-helix that typically repositions AAA+ large and small domains, in DNAHX it serves a distinct role by blocking the CTD swing during dynein’s mechanochemical cycle. This specificity suggests that the PS2i insertion in DNAHX is unique to the dynein family. Overall, the novel PS2i insertion in DNAHX broadens our understanding of AAA+ protein functional diversity.

### DNHD1 Maybe a Structural Ortholog of DNAHX

The dynein gene DNHD1, traditionally annotated as an “orphan” DHC gene, was previously thought to be a vestige of early DHC duplication with no functional role. However, recent genetic and functional studies suggest that DNHD1 is critical for ciliogenesis in both sperm cells and embryonic nodal cells [24, 25, 26]. Interestingly, AlphaFold3 predictions for the DNHD1 motor domain reveal structural features strikingly similar to DNAHX.

First, DNHD1 contains multiple disordered insertions throughout its motor domain, although not at the same sites as DNAHX (**Supplementary Fig. 5a, b**). Additionally, sequence analysis indicates that its AAA1 site is poorly conserved, lacking the catalytic Walker-B residues necessary for ATP hydrolysis (**Supplementary Fig. 5c**). These features imply that DNHD1, like DNAHX, may lack microtubule-based motility. Despite their lack of sequence homology, DNHD1 and DNAHX may both belong to a class of non-motile dyneins that evolved along distinct evolutionary pathways. Future biochemical and structural studies of DNHD1 and DNAHX will be essential to confirm their structural and functional relationship and to explore their roles in cilia assembly and regulation.

### Potential Role of DNAHX and Dynein-X in Sperm Flagellar Growth

Permanently locked in a pre-powerstroke state, DNAHX is structurally suited for forming phi particles, as the motor domain in phi dynein must remain in this state. This distinguishes DNAHX from other DHCs, which undergo an AAA1-dependent mechanochemical cycle that restricts many conformations incompatible with motor domain dimerization. Dyneins in the phi particle state are known to be autoinhibited, exhibiting low affinity for microtubules—a critical regulatory mechanism that prevents unneeded dynein activation [44]. This inhibition is essential for dyneins operating within specialized cellular structures like cilia or flagella, especially for cytoplasmic dynein-2 and axonemal dyneins. Phi particles of dynein-2 [43] and phi like conformations of OADs [46] have been shown to play a dual role: (1) inhibiting premature binding and activation on cytoplasmic microtubules and (2) promoting the compact conformation needed for docking onto intraflagellar transport (IFT) trains for ciliary delivery.

Most axonemal dyneins, however, are single-headed, and their inhibition and transport to the ciliary tip for activation remain unclear. This regulation is especially important in sperm cells, where flagella, often 50–150 *µ*m in length, significantly exceed typical cilia (<10 *µ*m). The extended length necessitates precise regulatory control to ensure correct flagellar growth and motility. DNAHX, a non-motile DHC identified in sea urchin sperm, could serve as a regulatory component for inhibiting and transporting axonemal dyneins during ciliogenesis. Specifically, DNAHX can serve as a ‘phi template’ to dimerize with other axonemal DHCs, forming a larger dynein-X complex.

We propose the following model for DNAHX and dynein-X’s role in sperm cells maturation (**Fig. 6f**). Although mature sperm cells no longer require IFT [48], IFT machinery is involved for axoneme assembly during spermatogenesis [48, 49, 50]. Near the basal body, a dynein complex pool—including dynein-2, OADs, IADs, and DNAHX—exists in a primed but inhibited state. Dynein-2 and OAD form phi particles to prevent premature cytosolic activation. DNAHX, inherently locked in a pre-powerstroke state, can form heterodimeric phi particle dynein with single-headed dyneins, inhibiting the activity of IADs. This phi particle conformation also facilitates dynein-X docking onto anterograde IFT trains. Once at the ciliary tip, the phi particle dynein-X transitions to open conformation, enabling IAD activation and docking onto DMTs, while DNAHX returns to recycle and package additional DHCs.

## Material and methods

### Sperm axonemal dynein purification from sea urchins

Isolation of sperm cells from sea urchins was conducted based on previous published protocol [51]. Live green sea urchins (*Lytechinus variegatus*) were purchased from gulf specimen marine lab (https://gulfspecimen.org/). For each batch, we used 12-16 sea urchins. Spawning was induced by the injection of 1ml of 0.5 M KCl into the perivisceral cavity and semen were collected.

The flagellar axoneme isolation and dynein purification were modified based on published protocols [52, 53, 54, 55]. Briefly, the semen was resuspended in isolation buffer (100mM NaCl, 5 mM imidazole/HCl, pH 7.0, 4 mM MgSO_4_, 1 mM DTT, 5 mM 2-mercaptoethanol, 0.2 mM PMSF) containing 20% sucrose. The suspension was lysed using a 15 mL Dounce tissue grinder (Wheaton) with 10-15 strokes on ice. The sperm head was pelleted twice via centrifugation at 3000g for 7 minutes. The demembraned axoneme was pelleted at 27,000g for 15 minutes. The pellet was washed twice and resuspended in isolation buffer without sucrose.

For dynein purification, 100 mg axoneme pellet was resuspended in 10mL isolation buffer with adjusted 0.6 M NaCl high salt and 1 mM ATP (10 mg/ml in 10 mL buffer). After incubation on ice for 30 minutes, the microtubule was pelleted at 27,000g for 20 minutes and the supernatant containing dyneins was collected and concentrated using a pre-equilibrated 100K Amicon Ultra-15 concentrator (Millipore). 1 mL concentrated supernatant was laid over the 5–25% linear sucrose gradient in dynein buffer (50 mM HEPES pH 7.4, 100 mM NaCl, 1 mM DTT, 0.1 mM ATP) and centrifuged at 153,000× g for 16 h at 4 °C in a swinging bucket rotor (a 16.8 mL SW32 Beckman polyallomer tube). The gradient was fractionated into 0.5 mL aliquots and the fractions containing OAD or IAD were determined by SDS-PAGE and negative staining microscopy. The fractions containing the two-headed dynein were pooled together and dialyzed against the dynein buffer for 6 h to remove the sucrose. The final dynein sample was concentrated to 5 mg/ml and immediately used for cryo-EM sample preparation.

### Negative-stain electron microscopy and 2D classification analysis

Dynein was diluted to 0.01-0.03 mg/ml in dynein buffer and 4*µ*L sample was applied to glow-discharged grids (300-mesh copper with carbon support, Electron Microscopy Sciences). The grid was manually blotting with a filter paper and stained with 2% uranyl acetate for 1 minute. The grids were then imaged using a Talos L120C microscope (Thermo Fisher Scientific) and a small dataset was collected to access the quality of the purified dyneins. The images were imported to cryoSPARC [56] for particle picking and 2D classification.

### Mass spectrometry

Mass spectrometry (MS) on the isolated dynein sample was performed at Keck Biotechnology Resource Laboratory, Yale University. The protein relative abundance was calculated in Scaf-fold 5 software (http://www.proteomesoftware.com/products/scaffold-5).

### Cryo-EM sample preparation and data collection

Dynein was diluted to 2 mg/ml and ATP concentration was adjusted to 3 mM for cryo-EM sample vitrification. 3*µ*L dynein sample was applied to Quantifoil holy carbon grids (R2/1, 300 mesh gold or R2/1, 400 mesh) and incubated in the chamber of the Vitrobot Mark IV unit (FEI) for 5 seconds at 4 °C and 100% humidity. Subsequently, the grids were blotted for 3-5 seconds and plunged into liquid ethane. The grids were then screened, and data were collected on a 200 keV Glacios electron microscope (Thermo Fisher Scientific) with a K3 direct detection camera (Gatan) with a magnification of 45000x, physical pixel size of 0.868 Å and total dose of 40 e-/Å2. In total, 7, 816 movies at a defocus from -1.2 *µ*m to -3.0 *µ*m were acquired and data collection was automated using SerialEM [57].

### Cryo-EM image processing

Preprocessing steps, including motion correction, CTF estimation, and particle picking, were conducted either in cryoSPARC Live [56] or via an in-house script utilizing MotionCor2 [58], GCTF [59], and Gautomatch. Cryo-EM scripts for real-time data transfer and on-the-fly preprocessing are available for download at https://github.com/JackZhang-Lab. The following reconstruction steps were conducted in cryoSPARC [56].

The initial particles were picked using a blob picker (dimension: 140Å, covering a single dynein motor domain) and extracted at a box size of 128 at 2.604 Å pixel size (bin 3). The particles were subjected to several rounds of 2D classification to obtain high-quality class averages of dynein motor domain. Ab initio reconstruction (6 classes) followed by heterogeneous refinement of these particles reveal medium resolution (5-8 Å) features of motor domain in pre and post powerstrokes. These initial reconstructions were then served as references for the following “cross-classification” scheme described previously in our work [60, 61]. The steps are summarized below:

The particles were re-picked using template-matching of the high-quality 2D class-averages. The cross-correlation score was set to low (e.g. 0.1) to enable sampling of all possible motors during the picking step. In total, this yielded 6,644,359 initial particles. These particles were extracted at bin 3 and then divided into 4 sets with each set containing around 1,500,000 particles. Each set was directly subjected to heterogeneous refinement (10 classes) using the initial references obtained before. This typically yielded 3-4 good classes with distinct features of motor domain while the other classes appeared as junk volumes. Due to the oversampled picking scheme, many good particles were hidden inside theses classes as well. To classify good particles from these junk classes, we merged the good classes and treat them as seed particles. We then divided the original particles into smaller sets with each set containing 500,000 particles. These particles, together with seed particles were merged in a heterogeneous refinement job. The seed particles provided stability of high-quality reconstructions during each iteration and was able to pool out more good particles from the junk classes.

After cross-classification, we separated the highquality motor domain reconstructions into two classes: pre-powerstroke state with a bent linker and post-powerstroke state with a straight linker and reextracted particles at a box size of 256 at 1.3 Å pixel size. A high-resolution heterogeneous refinement job (setting the “box size” parameter to 256) was then executed. The high-resolution information allowed the clear separation of motor domains at different conformations from different dynein heavy chains, with each class achieving 3.5-4.0 Å resolution at the stage of heterogeneous refinement. Each class was then refined to 2.8-3.3 Å resolution after homogeneous refinement, two rounds of global, local CTF refinement and local refinement.

To find potential interactor of DNAHX, we reextracted DNAHX particles using a larger box size of 256 at 2.6 Å pixel size (dimension: 666Å) and performed reference-free 2D classification. This revealed that a subset of DNAHX was forming a phi particle with another DHC. 3D reconstruction of this phi particle yielded a 10Å map.

### Protein identification, model building and refinement

To identify the corresponding DHC gene for each high-resolution map, we utilized recently developed AI-based automatic model tools, deepTracer [31] and ModelAngelo [30], to place the best-guessed amino acids into cryo-EM maps. This typically yield several polypeptide chains of each motor domain. We then extracted the longest chain and used “blastp” to search the closed dynein gene in in *Lytechinus variegatus* NCBI database. All maps had a single top hit during this step owing to the high-resolution map and long sequence of motor domain.

Following protein identification, ModelAngelo [30] was used to automatic build the models with sequence information. However, some regions with low local resolution were too flexible to be automatic built. To address this, we used Alphafold2 [34] to predict the individual AAA+ domain and then fitted the domain into the automatic built models. A manual editing step was conducted in Chimerax [62] and Coot [63, 64] to merge different models, connect missing loops, etc. Namdinator [65], a molecular dynamics flexible fitting tool, was employed to further refine the model into the cryo-EM map.

All models underwent iterative refinement using Phenix real-space refinement 1.21rc1-5190 [66] and manual rebuilding in COOT. The quality of the refined models was assessed using MolProbity [67] integrated into Phenix.

### Phylogenetic and structural analysis of DNAHX with canonical DHC genes

The canonical DHC genes from *Homo sapiens, Lytechinus variegatus, Ashbya gossypii, Caenorhabditis elegans, Chlamy reinhardtii, Danio rerio, Drosophila melanogaster, Kluyveromyces lactis, and Tetrahymena thermophila* were extracted from uniport or NCBI database using human DHC genes as references. The DNAHX from sea urchin gene was not found in these model organisms. The Phylogenetic analysis pipeline by ETE3 [68] was conducted using online server GenomeNet using default settings (https://www.genome.jp/tools-bin/ete). The tree was visualized in iTOL [69](https://itol.embl.de/).

To find the DNAHX orthologs, we blasted the sea urchin DNAHX’s gene against the NCBI database. The hits were manually curated such that each animal phylum has a presentative organism. To further confirm the found gene was true ortholog of DNHAX but not the other DHC genes, we blasted the gene to *Lytechinus variegatus* database to validate the top hit was indeed DNAHX. These DNAHX genes, along with canonical DHC genes were isolated and subjected to the same phylogenetic analysis pipeline. The multiple sequence alignment results were visualized in Jalview [70].

### Plots and molecular graphics

Plots were generated using GraphPad Prism. Molecular graphics were prepared using UCSF ChimeraX [62] and Pymol (https://www.pymol.org/).

## Data Availability

Models and cryo-EM maps have been deposited in the PDB and EMDB as follows: PDB-XXXX/EMDB-XXXXX for DNAHX motor domain, PDB-XXXX/EMDB-XXXXX for DNAH8 motor domain in pre-powerstroke state, PDB-XXXX/EMDB-XXXXX for DNAH8 motor domain in post-powerstroke state, EMDB-XXXXX for heterodimeric phi particle dynein.

## Acknowledgements

We thank the Zhang lab members for valuable discussions. We would like to thank K. Zhou, J. Lin, M. Llaguno, and S. Wu, for their help with cryo-EM data collection at Yale Cryo-EM facility. The Yale Cryo-EM Resource is funded in part by the NIH grant S10OD023603. We thank J. Wang at National Cancer Institute for the help of cryo-EM data collection. This research was, in part, supported by the National Cancer Institute’s National Cryo-EM Facility at the Frederick National Laboratory for Cancer Research under contract 75N91019D00024. We thank L. Wang, J. Kaminsky, and G. Hu at the Laboratory for BioMolecular Structure (LBMS) for help with cryo-EM data collection. The LBMS is supported by the DOE Office of Biological and Environmental Research (KP1607011). This work was funded by the NIH/NIGMS (GM142959 to K.Z.).

## Author Contributions

K.Z. designed the study. P.C., W.Z. and J.Y. isolated sea urchin sperms and purified proteins. P.C. prepared the cryo-EM samples, collected the data, processed the images, and built PDB models. P.C. and D.S.L. analyzed the data and generated figures. P.C. wrote the draft. P.C., D.S.L., and K.Z. edited and revised the manuscript. K.Z. acquired funding.

---

**Supplementary Figure 1:**
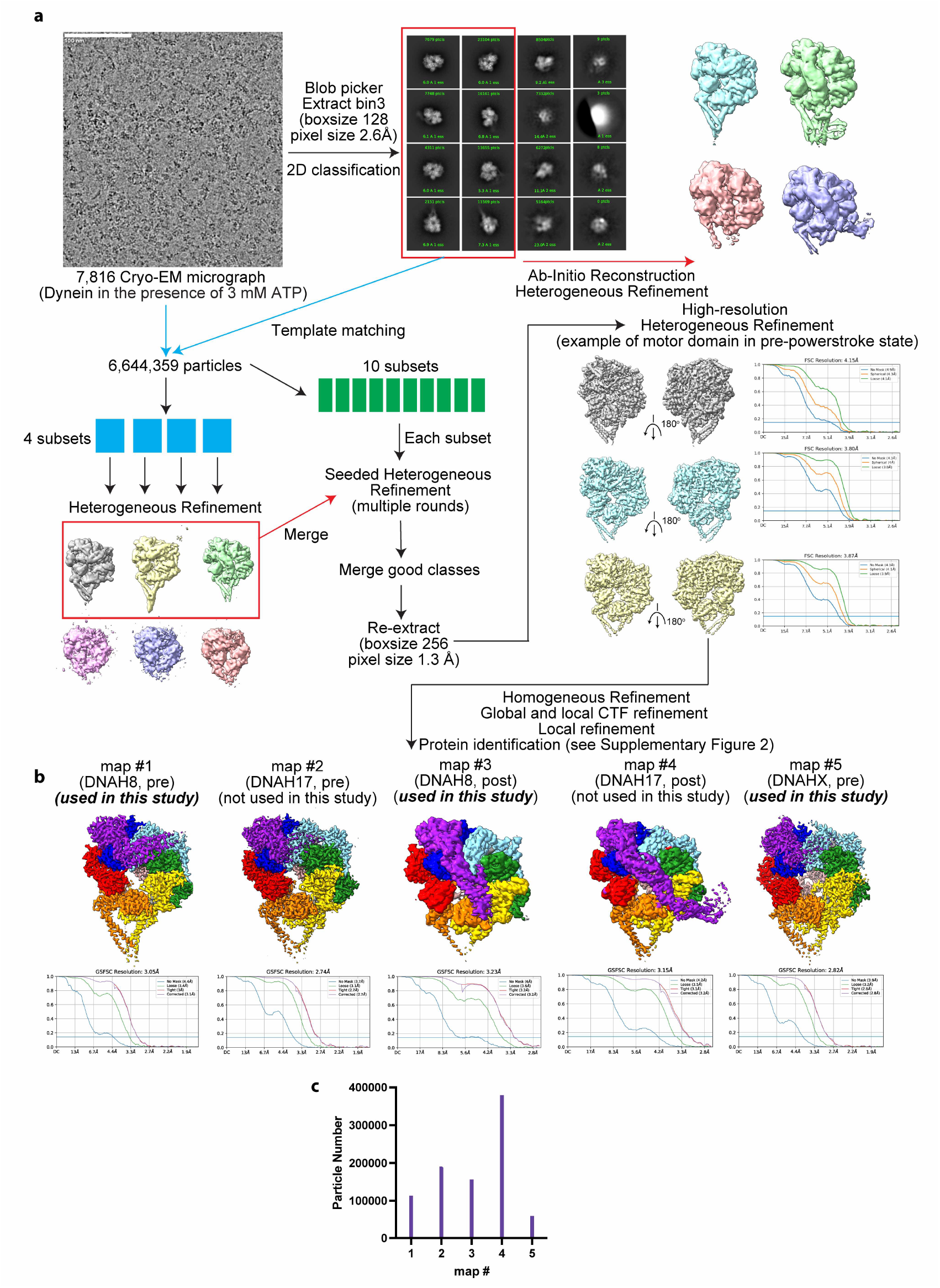
Cryo-EM data processing workflow. **(a)** Cross-classification workflow for cryo-EM image processing. **(b)** Final high-resolution reconstructions of motor domains from different DHC at different powerstroke states. **(c)** Particle number distribution of five high-resolution maps.

**Supplementary Figure 2:**
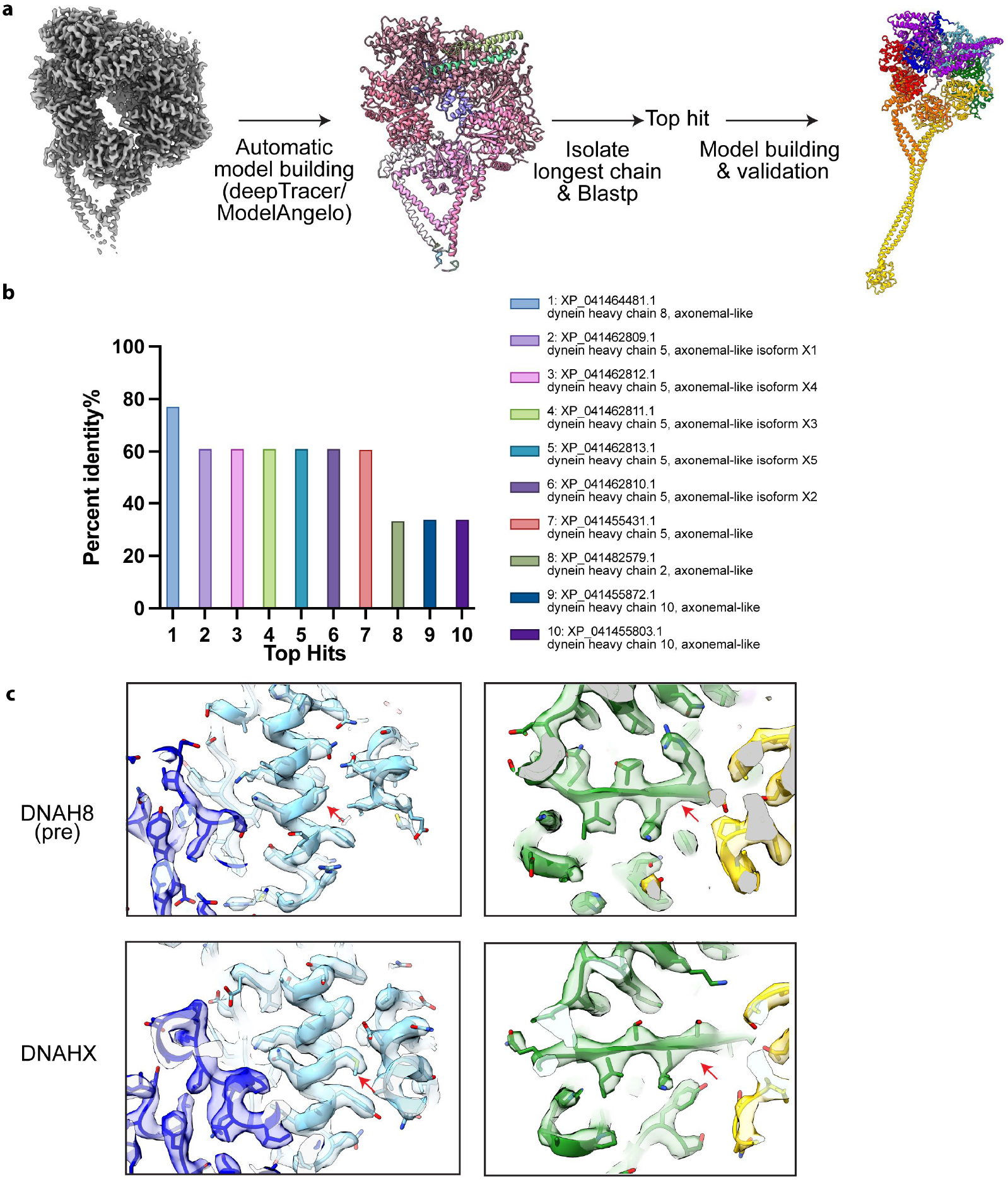
Protein identification workflow. **(a)** Automatic model building tools were used to build best-guessed model into cryo-EM maps. The protein sequence was isolated for Blastp search. **(b)** An example of search result. The top hit has a 77% percent identity while the second hit has a 61% percent identity. **(c)** Representative cryo-EM densities from DNAH8-pre and DHAHX. Similar regions are shown, and differences are highlighted using red arrows.

**Supplementary Figure 3:**
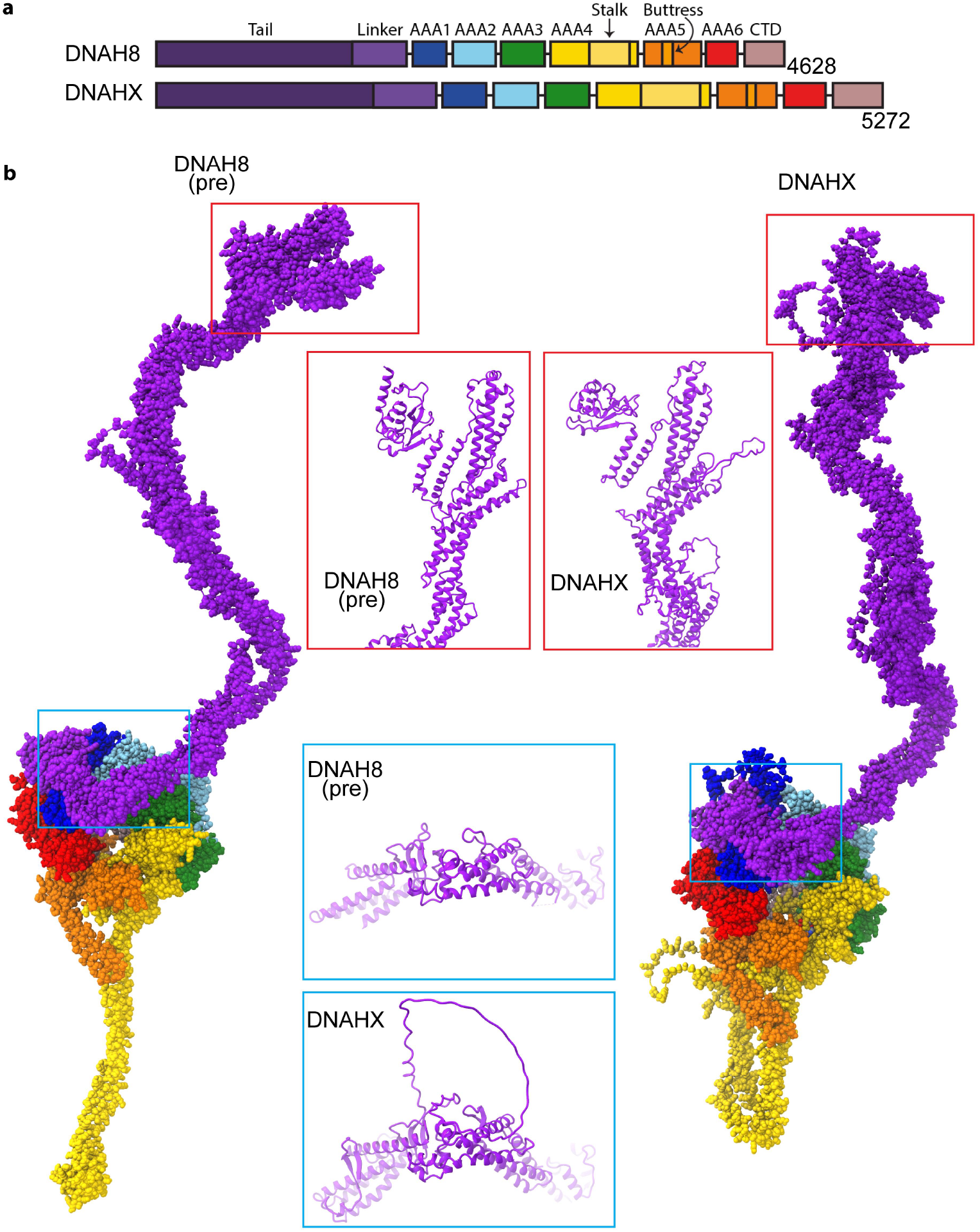
Structural comparison of full-length model of DNAHX and DNAHX8. **(a)** Domain diagram of DNAH8 and DHAHX. DNAHX doesn’t contain addition domains. The extended length is due to various insertions across the protein. **(b)** Full-length DHC models constructed from Alphafold2 predictions and cryo-EM maps. Two regions at the tail and linker regions are highlighted, showing the disordered insertions of DNAHX.

**Supplementary Figure 4:**
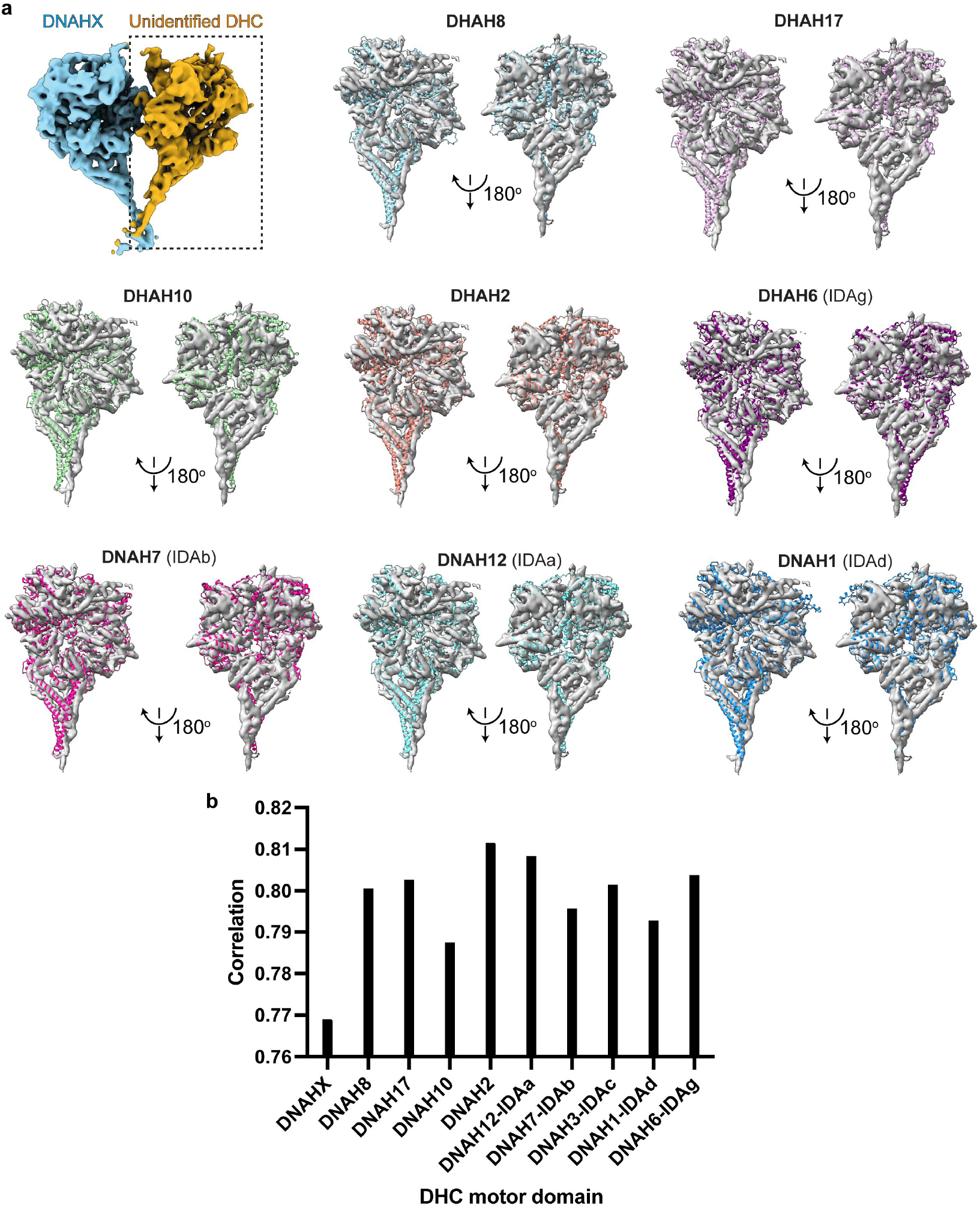
Structural fitting of DHC motor domains into cryo-EM density map. **(a)** Representative fitting of different DHC motor domains from sea urchin to density map. The pre-powerstroke models of motor domains are predicted by Alphafold3. **(b)** Correlation value among different fitted structures calculated by ChimeraX.

**Supplementary Figure 5:**
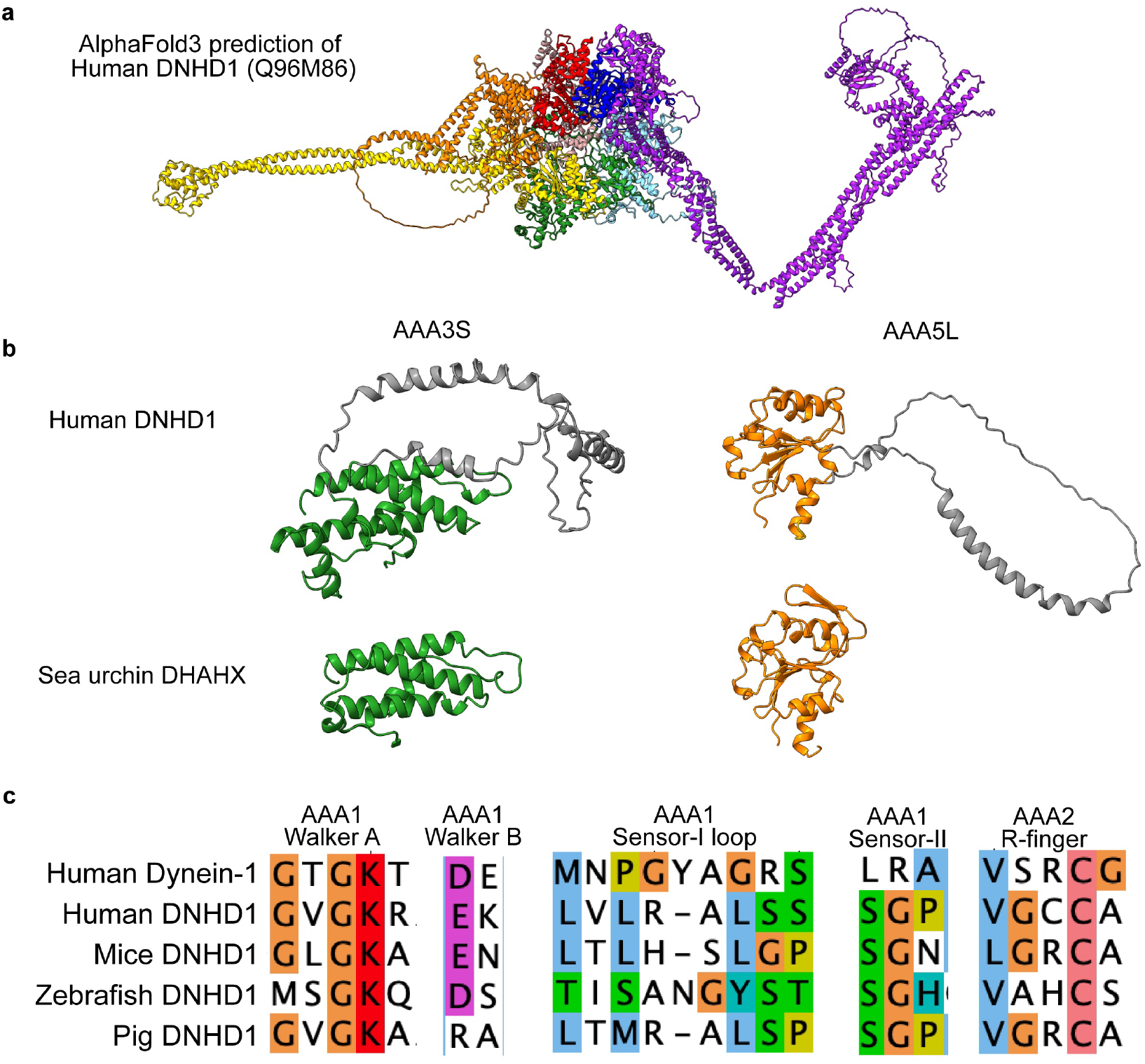
Structural features of DNHD1. **(a)** Alphafold3 predictions of human DNHD1 heavy chain. **(b)** Two examples of structural insertions of DNHD1. **c)** Alignment of key motifs at AAA1 sites among human dynein-1 and DNHD1 from various species.

**Supplementary Table 1:** Cryo-EM Data Collection, Refinement, and Validation Statistics.

| Native axonemal dyneins purified from sea urchin sperm |  |  |  |  |
| --- | --- | --- | --- | --- |
| Description | DNAH8<br>Pre-powerstroke | DNAH8<br>Post-powerstroke | DNAHX<br>Pre-powerstroke | DNAHX<br>Phi particle state |
| Facility | Yale ScienceHill-Cryo-EM facility |  |  |  |
| Microscope | Glacios |  |  |  |
| Voltage (kV) | 200 |  |  |  |
| Camera | K3 |  |  |  |
| Magnification | 45k |  |  |  |
| Pixel Size (Å) | 0.432 (super resolution) |  |  |  |
| Total Electron Exposure (e-/Å<sup>2</sup>) | 40 |  |  |  |
| Defocus Range (µm) | 1.5-2.7 |  |  |  |
| Symmetry Imposed | C1 | C1 | C2 | C1 |
| Num of mics | 7,816 |  |  |  |
| Initial Particles | 6,644,359 |  |  |  |
| Final Particles | 112,614 | 189,752 | 59,326 | 6,898 |
| Initial models | ab initio<br>AlphaFold2 | ab initio<br>AlphaFold2 | ab initio<br>AlphaFold2 | NA |
| Map pixel size | 0.868 | 1.302 | 0.868 | 1.736 |
| Map Resolution (Å) (FSC 0.143) | 3.0 | 3.3 | 2.8 | 7.9 |
| Map sharpening B-factor (Å<sup>2</sup>) | 80 | 80 | 66 | / |
| Model Composition |  |  |  |  |
| Non-hydrogen atoms | 22,862 | 23,609 | 24,117 | / |
| Protein residues | 2,847 | 2,934 | 2,998 | / |
| Ligands | MG:3<br>ATP:2<br>ADP:2 | MG:4<br>ATP:2<br>ADP:6 | MG:3<br>ATP:3<br>ADP:1 | / |
| Model vs. Data |  |  |  |  |
| FSC Map to Model (Å) (FSC 0.5) | 3.2 | 3.6 | 3.1 | / |
| Correlation coefficient (mask) | 0.89 | 0.79 | 0.91 | / |
| **B factors (Å<sup>2</sup>)** |  |  |  |  |
| Protein | 70.43 | 24.99 | 36.22 | / |
| Ligand | 52.71 | 14.96 | 22.49 | / |
| **R.m.s deviation** |  |  |  |  |
| Bond length (Å) | 0.003 | 0.003 | 0.003 | / |
| Bond angles (°) | 0.608 | 0.645 | 0.530 | / |
| **Validation** |  |  |  |  |
| Molprobity score | 1.48 | 1.86 | 1.46 | / |
| Clashscore | 8.41 | 10.25 | 7.71 | / |
| Rotamer outliers (%) | 1.07 | 0.00 | 0.15 | / |
| Ramachandran plot |  |  |  |  |
| Outliers (%) | 0.00 | 0.24 | 0.00 | / |
| Allowed (%) | 1.66 | 4.47 | 2.18 | / |
| Favored (%) | 98.34 | 95.29 | 97.82 | / |
| Rama-Z (whole) | 1.02 | 0.59 | 1.79 | / |

